# Ancient evolutionary origin of intrinsically disordered cancer risk regions

**DOI:** 10.1101/2020.06.15.152298

**Authors:** Mátyás Pajkos, András Zeke, Zsuzsanna Dosztányi

**Affiliations:** Department of Biochemistry, ELTE Eötvös Loránd University, Pázmány Péter stny 1/c, Budapest H-1117, Hungary; Research Centre for Natural Sciences, Magyar tudósok körútja 2, Budapest H-1117, Hungary

**Keywords:** intrinsically disordered regions, linear motifs, gene duplications, de novo, evolutionary origin

## Abstract

Cancer is a heterogeneous genetic disease that alters the proper functioning of proteins involved in key regulatory processes such as cell cycle, DNA repair, survival or apoptosis. Mutations often accumulate in hot-spots regions, highlighting critical functional modules within these proteins that need to be altered, amplified or abolished for tumor formation. Recent evidence suggests that these mutational hotspots can not only correspond to globular domains but also to intrinsically disordered regions (IDRs), which play a significant role in a subset of cancer types. IDRs have distinct functional properties that originate from their inherent flexibility. Generally, they correspond to more recent evolutionary inventions and show larger sequence variations across species. In this work we analyzed the evolutionary origin of disordered regions that are specifically targeted in cancer. Surprisingly, the majority of these disordered cancer risk regions showed remarkable conservation with ancient evolutionary origin, stemming from the earliest multicellular animals or even beyond. Nevertheless, we encountered several examples, where the mutated region emerged at a later stage compared to the origin of the gene family. We also showed the cancer risk regions become quickly fixated after their emergence, but evolution continues to tinker with their genes with novel regulatory elements introduced even at the level of humans. Our concise analysis provides a much clearer picture of the emergence of key regulatory elements in proteins and highlights the importance of taking into account the modular organisation of proteins for the analyses of evolutionary origin.

## INTRODUCTION

Most human genes are thought to have an extensive and very deep evolutionary history. In line with the thought “Nature is a tinkerer, not an inventor” [1] major human gene families date back to the earliest Eukaryotic evolutionary events, or even beyond. The very oldest layer of human genes encode metabolically, structurally or otherwise essential proteins that typically go back to unicellular evolutionary stages. Mutations to this core biochemical apparatus can prove disruptive to all aspects of cellular life, and indeed there are known mutational targets associated with genome stability and cancer. In contrast to these “caretaker” genes, a more novel set of genes have emerged at the transition to a multicellular stage. These “gatekeeper” proteins are involved in cell-to-cell communication, especially in early embryonic development and tissue regeneration. Gatekeeper genes that control cell division are among the best known cancer-associated oncogenes and tumor suppressors [2].

In order to establish the evolutionary origins of cancer genes, Domazet-Loso and Tautz carried out a systematic analysis based on phylostratigraphic tracking [3]. By correlating the evolutionary origin of genes with particular macroevolutionary transitions, they found that a major peak connected to the emergence of cancer genes corresponds to the level where multicellular animals have emerged. However, many cancer genes have more ancient origin and can be traced back to unicellular organisms. These trends seem to apply to the appearance of disease genes [4] and novel genes in general as well [5]. These studies were based on the evolutionary history of the founder domains. However, new genes can also be generated by duplication either in whole or from part of existing genes, when the duplicate copy of a gene becomes associated with a different phenotype to its paralogous partner. This mechanism can also influence the emergence of disease genes [5].

By taking advantage of the flux of cancer genome data, several new proteins have been directly identified to play a direct role in driving tumorigenesis during recent years [6]. One of the key signatures of cancer drivers is presence of mutation hotspot regions, where many different patients might show a similarly recurrent pattern of mutations [7]. These hotspots are typically located within well-folded, structured domains. However, many cancer associated proteins have complex modular architecture which incorporates not only globular domains but also intrinsically disordered segments, which can also be sites of cancer mutations. In our recent work, we systematically collected disordered regions that are directly targeted by cancer mutations [8]. While only a relatively small subset of such cancer drivers were identified, their mutations can be the main driver event in certain cancer types. These disordered regions can function in a variety of ways which include post-transcriptional modification sites, linear motifs, linkers, and larger sized functional modules typically involved in binding to macromolecular complexes [8]. In general, due to the lack of structural constraints, disordered segments are evolutionarily more variable [9]. In particular, linear motifs can easily emerge to a previously non-functional region of protein sequence by only a few mutations, or disappear as easily, leaving little trace after millions or billions of years [10]. However, elements fulfilling a critical regulatory function might linger on for a longer time. So far, the evolutionary origin of intrinsically disordered regions that have a critical function proven by a human disease association has not been analyzed.

In the current study, we studied the evolutionary origin of disordered cancer risk regions. For this, we used a dataset of cancer driving proteins in which cancer mutations specifically targeted intrinsically disordered regions [8]. We retrieved phylogeny data from the ENSEMBL Compara database. Using a novel conservation and phylogenetic-based strategy we determined the evolutionary origin not only at the gene level but also at the region level. In addition, we also investigated the emergence mechanism of disordered cancer risk regions and how evolutionary constraints, selection and gene duplications events influenced the fate of these examples. Finally, we presented interesting case studies that demonstrate the ancient evolutionary origin of these examples and the continuing evolution of their genes built around the critical conserved functional module.

## RESULTS

### Evolutionary origin of genes and regions

Altogether, we collected 36 cancer risk regions of 32 disordered proteins and investigated the evolutionary origin at the level of genes and regions. The age estimation of disordered cancer genes was obtained using the last common ancestor of descendents using the ENSEMBL supertrees, which includes phylogeny of gene families returning not only individual gene history, but also relationships of ancient paralogs and their history (see Material and Methods). Using this strategy instead of analysing the evolution of individual genes or simply the emergence of the founder domain, we could define the origin of regions more precisely, even the ancient ones, without introducing any bias of overprediction of origins. However, some ambiguity still remained and were manually checked (Supplementary Materials 1). The genes were traced back to Opisthokonta (in accordance with the ENSEMBL database) and divided into four major phylostratigraphic groups, which are associated with the emergence of unicellular, multicellular organisms, vertebrates and mammals.

In total agreement with previous results, we found that 21 disordered cancer proteins, the majority of cases, have emerged at the level of eumetazoa. 14 cases were found to be even more ancient and could be traced back to single cell organisms, at least to Opisthokonta. The only protein that emerged more recently, at the level of Vertebrates, was CD79B, the B-cell antigen receptor complex-associated protein β chain. Its appearance is in agreement with the birth of many immune receptors [11] and is assumed to be driven by the insertion of transposable elements.

In around half of the cases (21), the emergence of the mutated region was the same as the emergence of the protein. Strikingly, these included five cases (EIF1AX, HIST1H3B, MLH1, RPS15, SMARCB1) where not only the gene/protein but also the region primarily mutated in human cancers were very ancient and could be traced back to unicellular organisms. 15 regions with Eumetazoa and 1 with Vertebrata origin could be traced back to the same level as their corresponding gene. However, in several cases the emergence of the region was a more recent event compared to the emergence of the gene. Of these, 8 regions emerged at the Eumetazoa and 7 at the Vertebrate level. In general, there was only one level difference between the emergence of the gene and the region at this resolution. The only exception was SETBP1. In this case, the region itself emerged at the Vertebrate level. However, the gene could be traced back to Opisthokonta level, although the Eumetazoa origin cannot be completely ruled out (see Supplementary Materials 1). Overall, many of the disordered regions were more recent evolutionary inventions compared to the origin of their genes, and date back to the common ancestors of Eumetazoans or Vertebrates. Nevertheless, the ancestors of all of the regions were already present from the Vertebrate level.

### Position conservation

Overall, these results point to the ancient evolutionary origin of disordered regions involved in cancer, not only at the gene level, but also at the region level. To take a closer look, we also calculated the conservation of individual positions within the regions based both in terms of homologous substitutions and identity. Results show that these residues are highly conserved. 86% of the regions have more than 0.8 average conservation value even based on identities. Among the cases with the four lowest values, the conservation of VHL, CALR, and APC, which all correspond to relatively longer segments, was still relatively high. The only outlier was BCL2. In this case, the mutations are distributed along the N-terminal, encompassing the highly conserved BH4 motif but also the linker region between the BH4 and C-terminal part which is conserved only in mammals (Figure S1).

Next, we investigated how this average value is altered when only the highly mutated positions are considered. We repeated that analysis for positions that had at least 15 and 25 missense mutations, which slightly decreased the number of regions considered. The remaining 28 and 17 regions with positions having at least 15 and 25 mutations had 0.93, 0.89 and 0.96,0.92 average conservation values based on substitutions and identity, respectively (Figure 2C,D). This reflects a very clear trend with positions with a higher number of cancer mutations showing higher evolutionary conservation.

**Figure 1.**
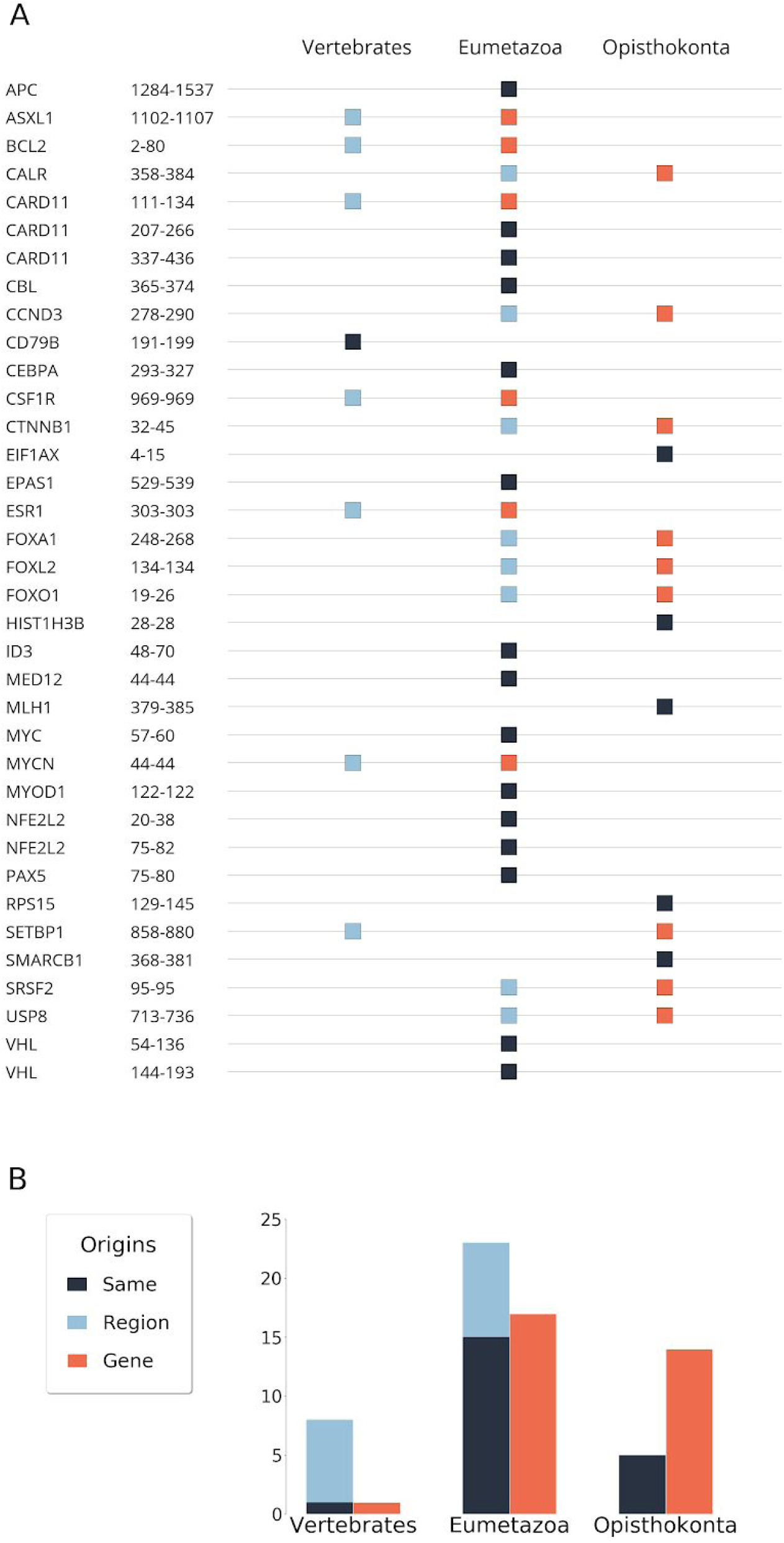
Conservation based evolutionary origin of disordered cancer regions and genes. **(A)** The orange and sky blue squares represent the origin of regions and genes, respectively. Gunmetal squares indicate the same evolutionary origin at both region and gene levels. **(B)** Summary barchart of origins in the three gene-age categories.

**Figure 2.**
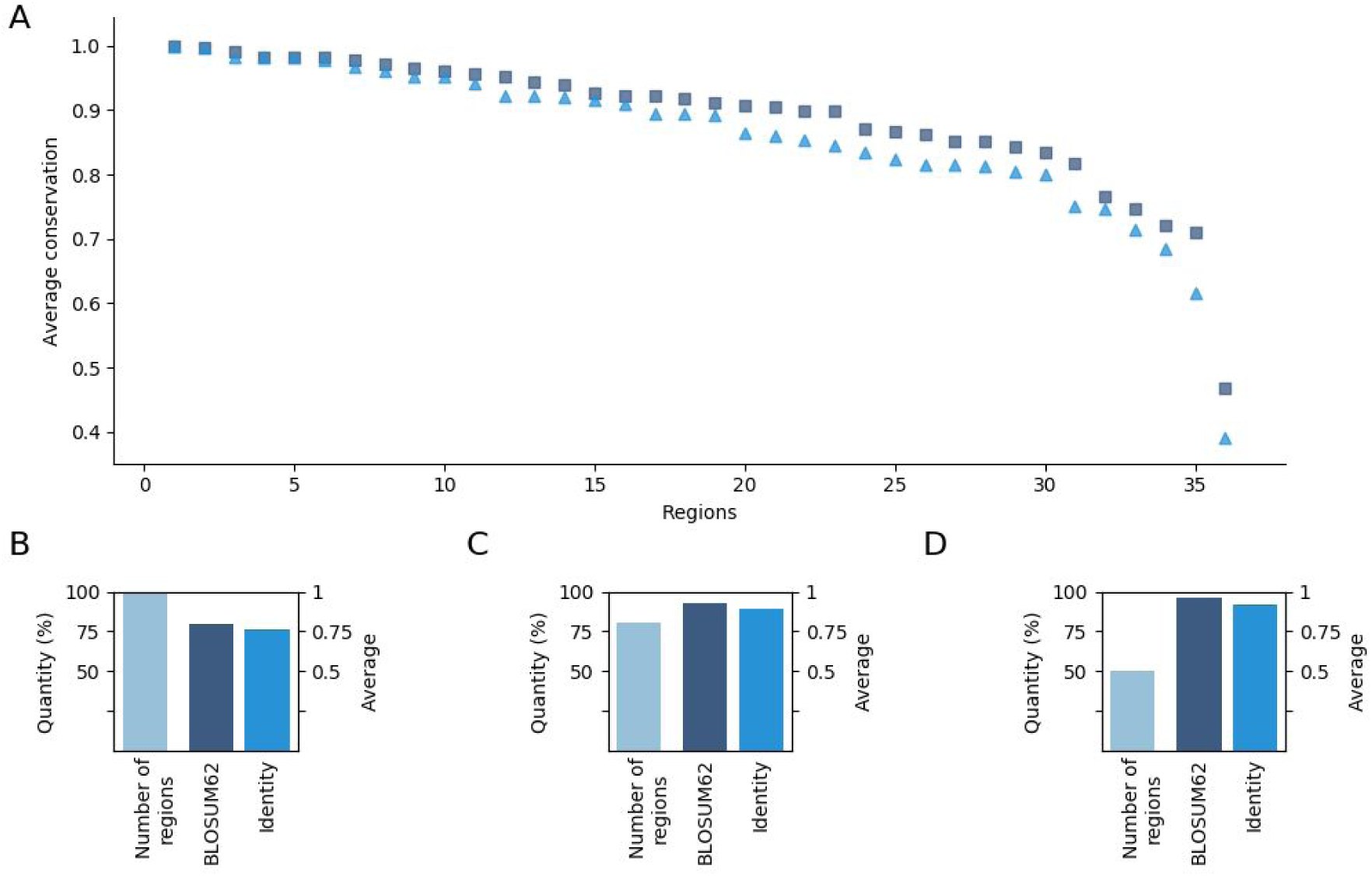
Representation of average conservation values of region positions having missense mutations. **(A)** Sorted average conservation values of each region having positions with at least one mutation. Squares and triangles represent BLOSUM62 and identity based conservation values, respectively. The outlier at the very end of the sequence is the region of BCL2. (**B-D**) Region number and average conservation value of regions having positions with at least 1, 15 and 25 mutations, respectively. The conservation values are based on BLOSUM62 and identity, and the number of regions are colored by dark, medium and sky blue, respectively.

We also collected sites of potential positive selection mapped onto our genes based on the Selectome database [12] which provides information on likely molecular selection both at the level of the evolutionary branch and the sequence position based on the ratio of non-synonymous and synonymous substitutions (ω). According to these results, positive selection affected only three genes on the human lineage, CALR, CTNNB1 and VHL in our dataset. All of these selections could be mapped onto the Vertebrates division with multiple positions (see Material and Methods) (Table 1).

**Table 1.** Positive selection within disordered cancer genes. Positions within cancer risk regions are colored blue. The numbers in brackets are the posterior probability of positive selection for each position.

| Gene | Positions under positive selection referring to the human protein sequence |
| --- | --- |
| CALR | **83**(0.971), **155**(0.971), **177**(0.990), **267**(0.995), **307**(0.994), **336**(0.991), **360**(0.999), |
| CTNNB1 | **121**(0.999), **206**(0.993), **250**(0.998), **287**(0.991), **411**(0.998), **433**(0.993), **525**(0.997), **552**(0.998), **556**(0.916) |
| VHL | **127**(0.957), **132**(0.942), **141**(0.923), **171**(0.947), **183**(0.963), **185**(0.920), |

However, these positions showed limited overlap with the mutated regions. In this case of CTNNB1 none of the positions under selection overlapped with the cancer mutated region, and there was only a single position in the case of CALR, which was not directly targeted by cancer mutations. In the case of VHL, 6 positions were detected with selective pressure and 5 of them were situated within the significantly mutated region. However, none of them corresponded to a highly mutated residue.

Taking advantage of an earlier analysis [13], we also analyzed if there was any human specific positive selection. As the ω based approach can not be used without uncertainty to identify human-specific positive selection, this work relied on the McDonald and Kreitman (MK) test, which compares the divergence to polymorphism data using closely related species, as human and chimp. There was only a single entry in our database, ESR1 that showed human specific evolutionary changes (see Case studies).

### Contribution of duplications to the emergence of disease risk regions

Gene duplications often drive the appearance of a novel function through the process called neofunctionalization. In these cases, after a duplication event one copy may acquire a novel, beneficial function that becomes preserved by natural selection. Here, we have evaluated whether the emergence of disordered cancer regions correspond to such neofunctionalization events. For this analysis, we collected paralog sequences and evaluated if there were regions present in these sequences that showed clear evolutionary similarity to the cancer mutated region.

The evolutionary history of many genes are quite complex and can involve multiple duplication events. We focused on the level where the cancer regions emerged and distinguished the following scenarios based on the relationship between the duplication and the presence of the region among the paralogs. The first scenario corresponds to duplication induced neofunctionalization. In this case, an ancient cancer region emerged directly after a given gene duplication and became preserved in only one of the branches that appeared after the duplication (Figure 3A). There are two basic scenarios when the duplication cannot be directly linked with the emergence of the regions. One possible scenario is when both branches contain the region, which indicates that the region must have emerged before the duplication (Figure 3B). The other possible scenario is when the region emerged at a later evolutionary stage after a duplication, and duplication cannot be directly linked to neofunctionalization (Figure 3B).

**Figure 3.**
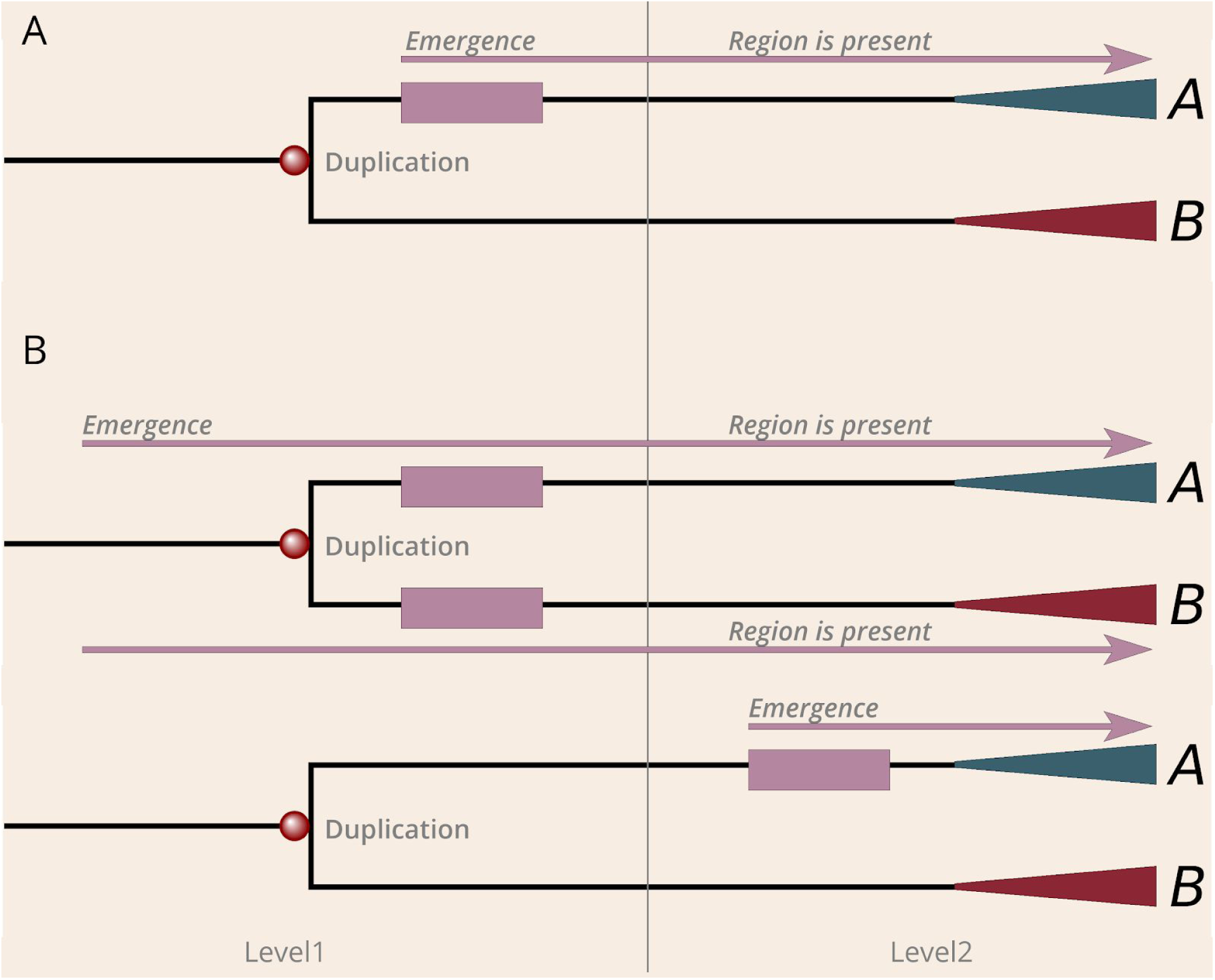
The mechanisms of emergence of regions by neofunctionalization and de novo. (**A**) Demonstration of the model of duplication induced (neofunctionalization) cancer region emergence. (**B**) Depiction of the two sub-scenarios of the de novo region emergence. Mallow boxes and arrows explains the evolution of the region. Red and green triangles symbolize the further evolution of paralogs after gene duplications.

Surprisingly, the duplication induced neofunctionalization was much less common than we expected, with only 7 cases showing this behaviour. One example for this scenario is presented by the β-catenin family, where the degron motif [14] based cancer risk region emerged after duplication is present only on the branch of β-catenin and junctional plakoglobin (JUP). In contrast, we found that 23 regions have evolved by de novo emergence, which seemed to be the dominant mechanisms for the emergence of the analyzed cancer mutated disordered regions (Figure 4A). For example, ID3 underwent multiple duplications, but all paralogs contain the cancer risk region, which indicates that the region emerged prior to the duplication. Another example is ESR1, in which case the paralogs were born at the level of Eumetazoa, however, this event is not directly linked to the emergence of the cancer region, which appeared only at the level of the ancient vertebrates. In addition, there were two singletons in our dataset, RPS15 and SMARCB1, which did not have any detectable paralogs. In the cases of ASXL1, CCND3, SETBP1 and the first region of CARD11, the evolutionary scenarios could not be unambiguously established. These six examples formed the Other group.

**Figure 4.**
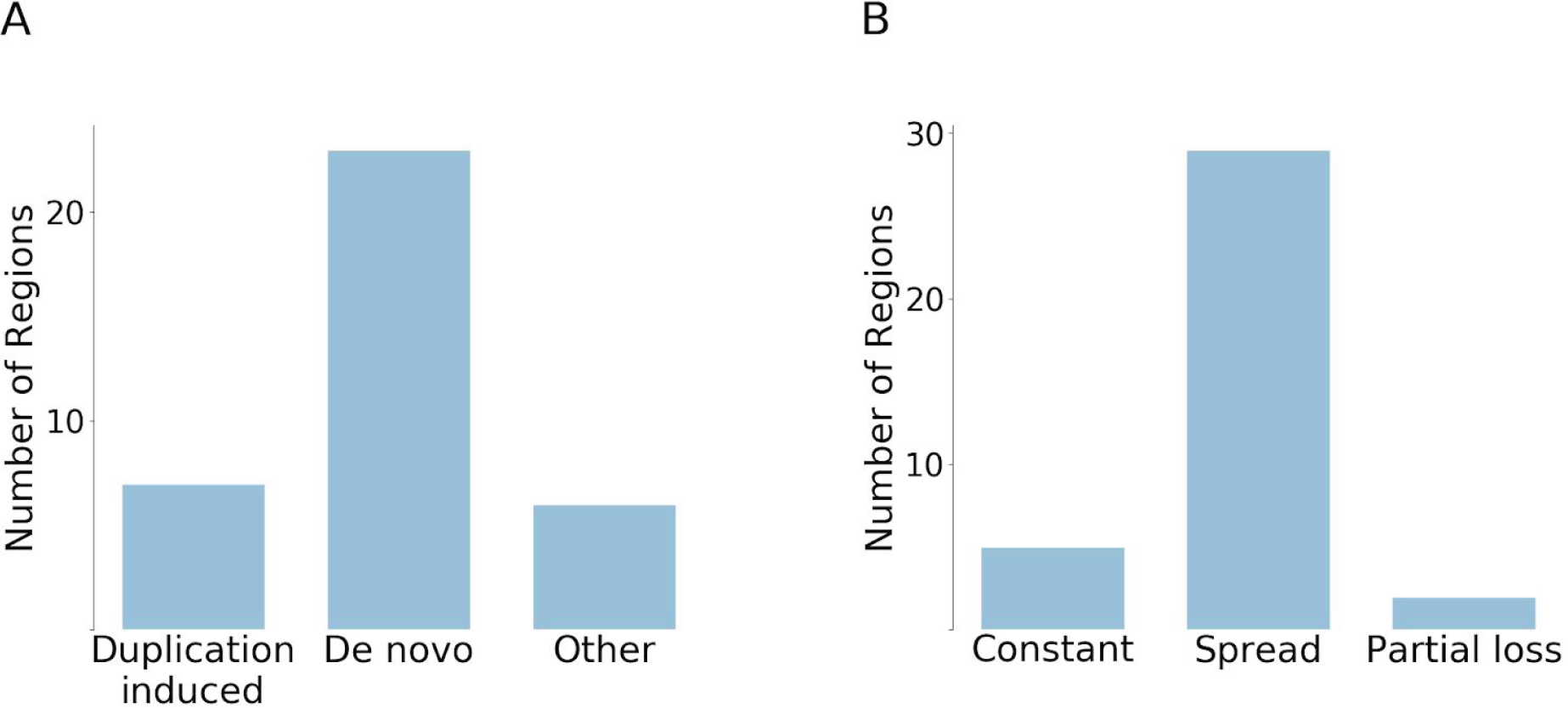
Categorization of emergence scenarios and evolutionary fates of cancer regions. **(A)** The number of regions that have emerged by duplication or de novo. 6 regions were not categorized (Other). **(B)** Classification of cancer regions in terms of their evolutionary fate after emergence.

We also analyzed if additional duplication events occurred after the emergence of regions and whether these novel paralogues retained the regions. There are basically three scenarios that can occur: (i) the region is preserved without any further duplications; (ii) the region spreads and becomes preserved in all of the novel duplicates; (iii) partial loss scenario, i.e. the region is preserved in some duplicates but is lost in others. Our results show that the most common evolutionary fate is the second one (Figure 4B). In 29 cases at least one duplication that inherited the region can be observed after the emergence of the cancer region. In contrast, only five regions were not duplicated. Some ancient cases, such as MLH1 and USP8, are also included among the non-duplicated ones, which means that the reason for the lack of duplications is not the short evolutionary time. The partial loss scenario was observed in only two cases, in the case of VHL and NFE2L2. For instance, in the case of VHL, there was a relatively recent gene duplication at the level of Mammals. While the N-terminal segment is present on both paralogs (VHL and VHLL), the C-terminal segment is only present in VHL but was lost from VHLL. In a similar fashion, NFE2L2 underwent a more recent gene duplication at the level of Vertebrates, but the newly emerged paralog did not retain the two linear motifs that are primarily targeted by cancer mutations.

### Case studies

#### MLH1

One of the most ancient examples in our dataset corresponds to MLH1, an essential protein in DNA mismatch repair (MMR). As one of the classic examples of a caretaker function, mutations of MLH1 can lead to cancer by increasing the rate of single-base substitutions and frameshift mutations [15]. Several positions of MLH1 are mutated in people with Lynch syndrome also known as hereditary nonpolyposis colorectal cancer (HNPCC). However, according to the COSMIC database of somatic cancer mutations, the most common mutation of MLH1 is V384D. Mutational studies of V384D using yeast assays and in vitro MMR assay did not indicate a strong phenotype, but still showed a limited decrease of MMR activity [16]. However, it was shown that the (mostly germline) V384D variant is clearly associated with increased colorectal cancer susceptibility [17] and it is highly prevalent in HER2-positive luminal B breast cancer [18].

MLH1 is an ancient protein that is present from bacteria to humans. It has a highly conserved domain organization that involves ordered N- and C-terminal domains connected by a disordered linker [19]. This underlines the functional importance not only of the structured domains but also of the connecting disordered region. In our previous work we identified the region from 379 to 385 to be significantly mutated [7], which is located within the disordered segment. Recently, it was shown that the linker can regulate both DNA interactions and enzymatic activities of neighboring structured domains [20]. In agreement with the linker function, both the composition and length of this IDR are critical for efficient MMR. Overall, most of the linker shows relatively low sequence conservation, however, the identified cancer risk region is highly conserved from across all eukaryotic sequences (Figure 5), in an island-like manner. Although the exact function of this region is not known, the strong evolutionary conservation indicates a highly important function, not yet explored in detail.

**Figure 5.**
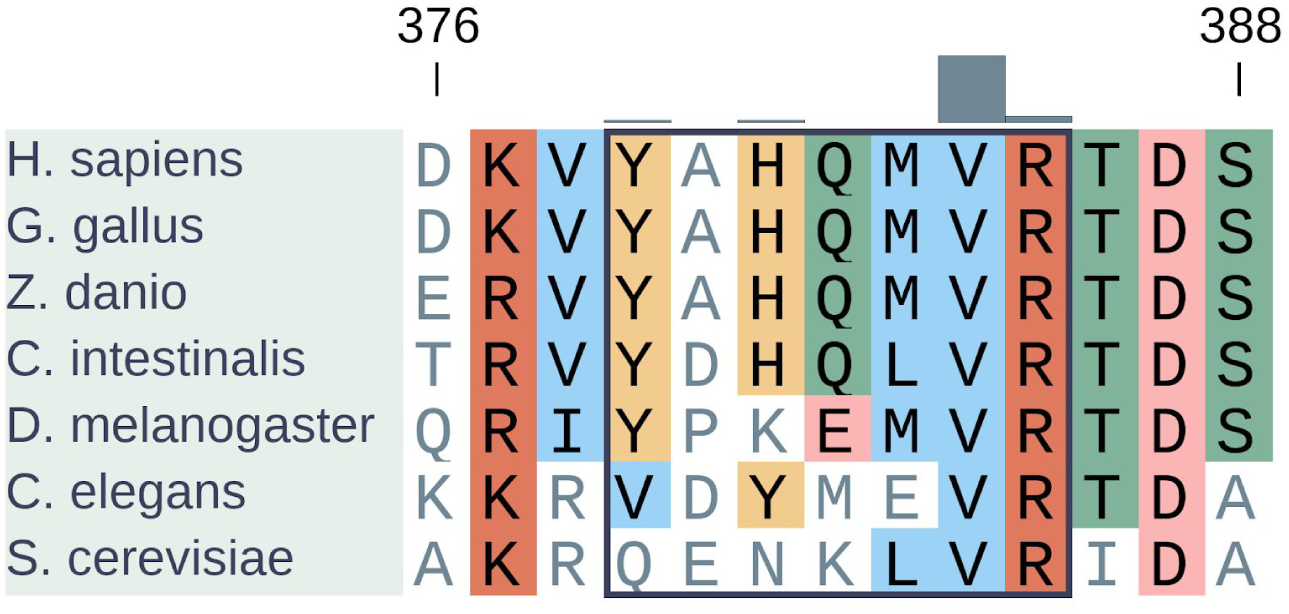
Segment of multiple sequence alignment of MLH1 generated with MAFFT [21]. Cancer region is indicated by a rectangle. Missense mutation distribution is depicted by gray bars.

#### VHL

VHL is a tumor suppressor protein that possesses an E3 ligase activity. It plays a key role in cellular oxygen sensing by targeting hypoxia-inducible factors for ubiquitylation and proteasomal degradation. To carry out its function, VHL forms a complex with elongin B, elongin C, and cullin-2 and the RING finger protein RBX1 [22,23]. VHL has an α-domain (also known as the VHL-box, residues 155 to 192) which forms the principal contacts with elongin C, and a larger β-domain (residues 63 to 154) which directly binds the proline hydroxylated substrate, HIF1α. The positions mutated across various types of cancers cover a large part of the protein, including both the α and β domains. While these regions form a well-defined structure in complex with elongin B, elongin C, and cullin-2, they are disordered in isolation and rapidly degraded [24].

The VHL gene emerged de novo at the level of Eumetazoa together with HIFα and PHD, the other key components of the hypoxia regulatory pathway. However, more recently, the gene underwent various evolutionary events. The VHL gene showed slightly higher evolutionary variations compared to other cancer risk regions (Figure 2). Some positions, including K171 showed signs of positive selection at the level of Sarcopterygii, which might implicate an important evolutionary event. It was shown that the SUMO E3 ligase PIASy interacts with VHL and induces VHL SUMOylation on lysine residue 171 [25]. VHL also undergoes ubiquitination on K171 (and K196), which is blocked by PIASy. In the proposed model of the dynamic regulation of VHL, the interaction of VHL with PIASy results in VHL nuclear localization, SUMOylation and stability for blocking ubiquitylation of VHL. Meanwhile, PIASy dissociation with VHL or attenuation of VHL SUMOylation facilitates VHL nuclear export, ubiquitylation and instability. This dynamic process of VHL with reversible modification acts in a concert to inhibit HIF1α [26].

A novel acidic repeat region appeared at the N-terminal region of the protein at the level of Sarcopterygii, and this region underwent further repeat expansion in the lineage leading up to humans (Figure 6). These GxEEx repeats are generally thought to confer additional regulation to the long isoform of VHL (translated from the first methionine), with a number of putative (USP7) or experimentally detected (p14ARF) interactors [27]. Although poorly studied, this repetitive region also seems to harbour casein kinase 2 (CK2) phosphorylation as well as proteolytic cleavage sites, regulating VHL half-life (consistent with a deubiquitinase, such as USP7 binding role) [28]. As a result of a recent gene duplication, the human genome even encodes a VHL-like protein (VHLL) which has lost the C-terminal segment including the α domain. Consequently, VHLL cannot nucleate the multiprotein E3 ubiquitin ligase complex. Instead, it was suggested that VHLL functions as a dominant-negative VHL to serve as a protector of HIF1α [29]. This example demonstrates that while the basic cancer risk region remains largely unchanged during evolution, additional regulatory mechanisms can emerge to further fine-tune the function of the protein.

**Figure 6.**
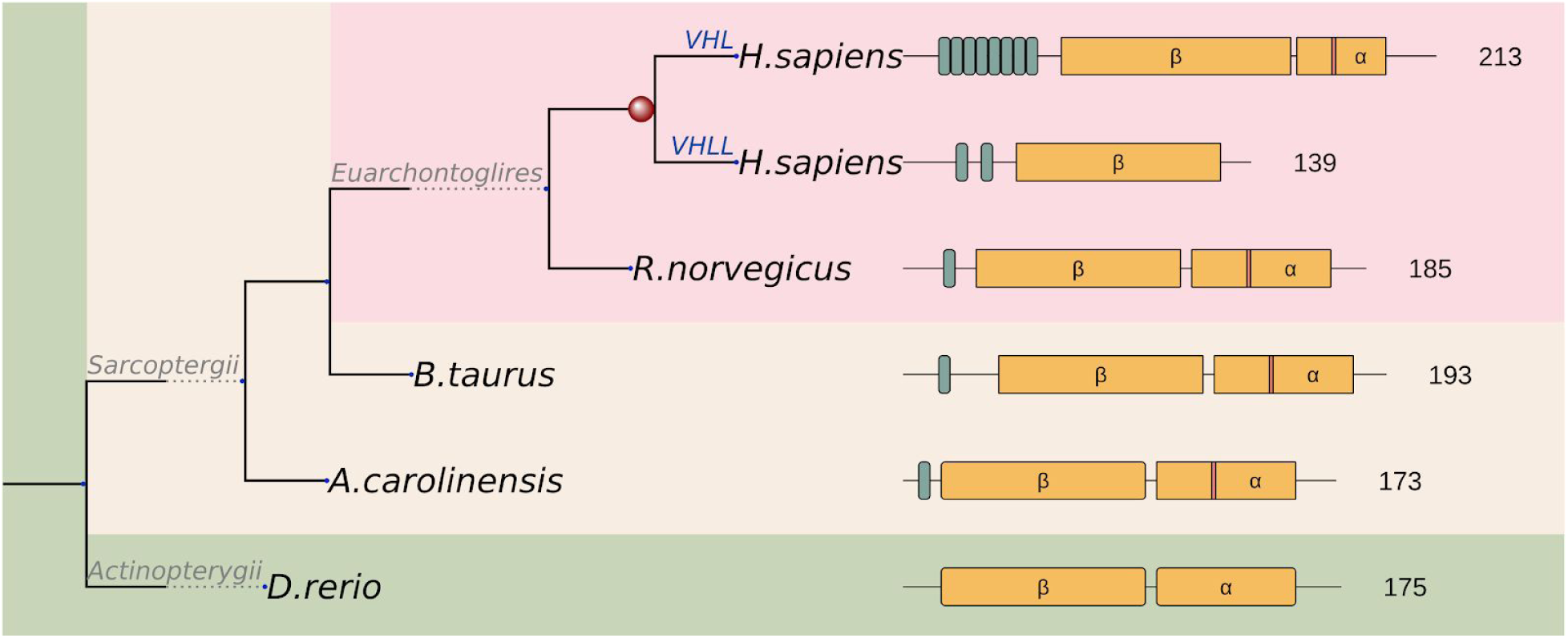
Schematic representation of the evolutionary scenario of the VHL family and their functional units. Repeat units in varying numbers and the α and β core domains are depicted by green and yellow boxes, respectively. Red stripe in the α domain of human VHL indicates K171 identified to emerge by positive selection on the Sarcopterygii branch (mapped K171 to other sarcopterygii are also indicated by red stripes).

#### ESR1

Estrogen receptor 1 (ESR1) is a member of the nuclear hormone receptor family with Eumetazoan origin. The most common mutation in both primary and tamoxifen therapy associated samples corresponds to a single mutation (K303R). This single site emerged more recently (Figure 7) and is located in a rather complex switch region adjacent to the ligand-binding domain (Figure S2). The highly mutated K303 of ESR1 (more than 200 K303R missense mutations are seen in COSMIC) is a part of a motif-based molecular switch region involving several mutually exclusive PTMs. At positions 302, 303 and 305 methylation by SET7/9, acetylation by p300 and phosphorylation by PKA or PAK1 were observed in previous studies, respectively [30–34]. Our results show that this region is conserved only in Sarcopterygii, which means a relatively young evolutionary origin of the switching mechanism. However, while the methylation and acetylation sites are well conserved, the phosphorylation motif appears to be specific only to *H. sapiens*. We came to this conclusion, because R300 and K302 as well as L306 are required for the protein kinase A (PKA) phosphorylation consensus and the oncogenic mutation K303R is expected to turn this region into an even better PKA substrate [35,36]. Curiously, these residues are not found in any other mammal, supposing species specific adaptive changes.

**Figure 7.**
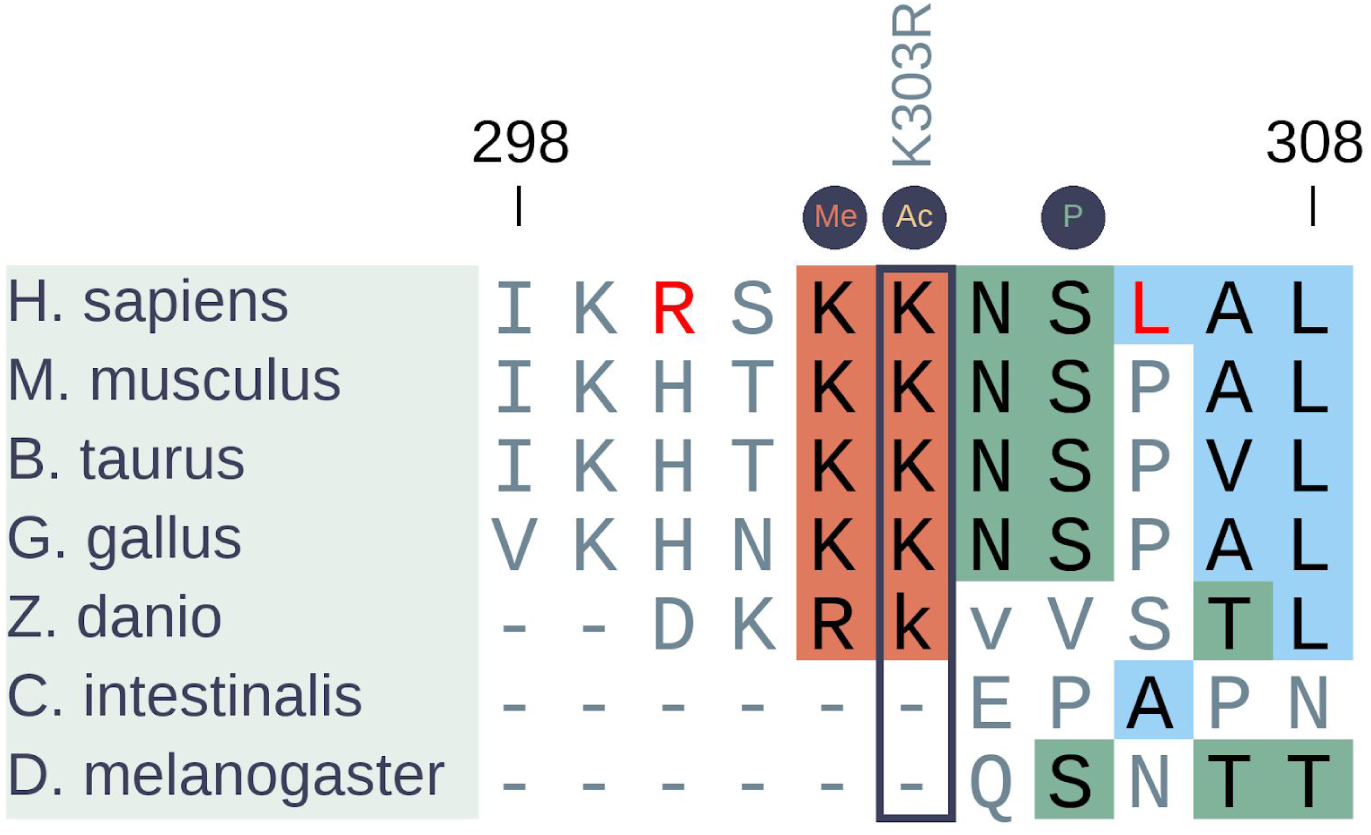
Insertion-free sequence alignment of the PTM site based cancer region of ESR1 generated with MAFFT [21]. Borders of non-depicted insertion of zebrafish are indicated by lower case letters. The highly mutated position (K303R) is highlighted by a rectangle. PTM sites are indicated by circles above the alignment. *H. sapiens* specific changes are colored by red.

Comparison of substitutions and polymorphic sites is a powerful approach to identify specific changes in a pair of closely related species, like *H. sapiens* and chimpanzee. Relying on this approach, 198 of 9785 analyzed genes were identified to show human-specific changes including ESR1 [13]. In ESR1 there are 3 more changes besides R300 and K306 (L44, Q502, S559) between *H. sapiens* and chimp that are also thought to be adaptive substitutions according to the MK test. Phosphorylation of S559 was experimentally identified suggesting this residue is also a *H. sapiens* specific PTM [37,38], but there is no specific data in the literature about the biological function of L44 and Q502. Yet we know that phosphorylation of S305 allows the increase of estrogen sensitivity by external stimuli other than steroids, and permits ESR1 activity even when the canonical estrogen effect is completely blocked by tamoxifen [34,36]. In mice, ESR1 activity is essential for estrogen effect and normal estrous episodes [39,40]. Although we lack information, we theorize that this human-specific signaling crosstalk might somehow be connected to the continuous menstrual cycle of *H. sapiens* (quite unusual among mammals), or some other human-specific reproductive adaptation.

## Discussion

In our study we aimed to estimate the evolutionary origin of disordered regions that are specifically targeted in cancer. Intrinsically disordered protein regions play essential roles in a wide-range of biological processes and can function as linear motifs, linkers or other intrinsically disordered domain-sized segments [41]. They are integral parts of many cancer associated proteins and in a smaller number of cases, they can also be the direct targets of cancer driving mutations. In general, IDRs are believed to be of more recent evolutionary origin, and exhibit higher rates of evolutionary variations compared to that of folded globular domains [9]. However, this is not what we see in the case of disordered cancer genes. Instead, we observed that cancer-targeted disordered regions are extremely conserved, with deep evolutionary origins which underlines their critical function. The two main ages for emergence of disordered cancer genes can be linked to unicellular organisms and the emergence of multicellularity, in agreement with the result of phylostratigraphic tracking of cancer genes in general [3].

One of the most unexpected findings of our study is the examples of disordered cancer genes that can be traced back to unicellular organisms. Mechanistically, the group of cancer genes that emerged in unicellular organisms were suggested to play a caretaker role and contribute to tumorigenesis by increasing mutation rates and genome instability. In contrast, cancer genes that emerged at the level of multicellularity were suggested to typically have a gatekeeper function and promote tumour progression directly by changing cell differentiation, growth and death rates [42]. MLH1 is one of the best characterized examples of a gene with a caretaker function [43]. It is involved in mismatch repair (MMR) of DNA bases that have been misincorporated during DNA replication. Thus, disruptive mutations of MLH1 greatly increase the rate of point mutations in genes and underline various inherited forms of cancer. However, the most commonly seen alterations in patients are located in the flexible internal linker. Mutational studies indicate that this highly conserved segment might not be directly involved in MMR but likely have an important, currently uncharacterized function. The other ancient examples are also involved in basic cellular processes, however they are associated with a broader set of functions. HIST1H3B, SMARCB1 and SETBP1 are involved in epigenetic regulation and their mutations can alter gene expression patterns [44,45]. Mutations of EIF1AX and RPS15 are likely to perturb translation events [46,47]. However, SRSF2, which is responsible for orchestrating splicing events, can also have a global influence on cellular states [48]. Therefore, the caretaker function is also a subject of evolution and some of its components emerged as a result of more recent evolutionary events.

A clear novelty of our approach is to focus at the origin of sub-gene elements: regulatory regions, modules and domains, instead of full genes. The genes can be built around founder genes that have an extremely ancient origin, but their biological function and regulation can change fundamentally during subsequent evolution. In several cases, the origin of the cancer mutated region was substantially more recent than the origin of the gene. Nevertheless, after their emergence disordered cancer regions got fixated rapidly and showed little variations afterwards. However, their evolution at the gene level was not set in stone and there are several indications that this process continues indefinitely. In several cases, the cancer genes underwent gene duplications, further regulatory regions were added, or fine-tuned by changing some of the less critical positions. We highlighted a fascinating case when such an event occured when our species, *H sapiens* separated from its primate relatives.

In general, the rate of gene duplications is very high (0.01 per gene per million years) over evolution, which provides the source of emergence of new evolutionary novelties [49]. According to the general view, paralogs go through a brief period of relaxed selection directly after duplications, which time ensures the acquisition of novelties, and subsequently experience strong purifying selection preserving the newly developed function. However, our results showed that only a few disordered cancer regions have emerged in a duplication induced manner and the vast majority of disordered cancer regions emerged by de novo, independent of duplications. The evolution of disordered regions is better described by the ex-nihilo motif theory, which is based on the rapid disappearance and emergence of linear motifs by the change of only a few residues within a given disordered protein segment [10]. This evolutionary phenomenon is commonly observed in the case of linear motifs, for example in the case of NFE2L2. This protein carries a pair of crucial linear motifs that have emerged in the ancient eumetazoa, but are not preserved in the most recent duplicates. In an evolutionary biology aspect, our results suggest that the evolution of functional novelties in the case of disordered region mediated functions requires a more complex model.

Exploring the evolutionary origin of cancer genes is an important step to understand how this disease can emerge. This knowledge can also have important implications of how their regulatory networks are disrupted during tumorigenesis and can be incorporated into developing improved treatment options [50]. In this work, we focused on a subset of cancer genes that belong to the class of intrinsic disordered proteins, that rely on their inherent flexibility to carry out their important functions. While the selected examples represent only a small subset of cancer genes, they are highly relevant for several specific cancer types [8]. In general, disordered proteins are evolutionarily more variable compared to globular proteins, however, the disordered cancer risk regions showed remarkable conservation with ancient evolutionary origin, highlighting their importance in core biological processes. Nevertheless, we found several examples, where the region specifically targeted by cancer mutations emerged at a later stage compared to the origin of the gene family. Our results highlight the importance of taking into account the complex modular architecture of cancer genes in order to get a more complete understanding of their evolutionary origin.

## Materials & Methods

### Dataset

We used a subset of the previously identified disordered cancer risk regions [8]. These regions were identified based on genetic variations collected from the COSMIC database [51] using the method that located specific regions that are enriched in cancer mutations [7]. Mapping was not feasible for CDKN2A isoform (this isoform was not found in the ENSEMBL database we used in our study), hence this protein was excluded from the further analyses. Proteins, in which both disordered and ordered cancer regions were identified, were filtered out in order to be able to focus clearly on the disordered regions. Regions that were primarily mutated by in-frame insertion and deletion and contained less than 15 missense mutations were also excluded because of our conservation calculation method (see below). Finally, histone proteins were merged keeping the single entry of HIST1H3B. Ultimately, we obtained a list of 36 disordered cancer risk regions of 32 proteins (APC [1284-1537], ASXL1 [1102-1107], BCL2 [2-80], CALR [358384], CARD11 [111-134; 207-266; 337-436], CBL [365-374], CCND3 [278-290], CD79B [191-199], CEBPA [293-327], CSF1R [969-969], CTNNB1 [32-45], EIF1AX [4-15], EPAS1 [529-539], ESR1 [303-303], FOXA1 [248-268], FOXL2 [134-134], FOXO1 [19-26], HIST1H3B [28-28], ID3 [48-70], MED12 [44-44], MLH1 [379-385], MYC [57-60], MYCN [44-44], MYOD1 [122-122], NFE2L2 [20-38; 75-82], PAX5 [75-80], RPS15 [129-145], SETBP1 [858-880], SMARCB1 [368-381], SRSF2 [95-95], USP8 [713-736], VHL [54-136; 144-193]).

### Evolutionary framework

In this work we calculated the evolutionary origin of cancer risk regions within our dataset of disordered proteins. Our approach focused on the age of orthologous gene families, instead of focusing on the evolutionary origin of founder domains. Assignment of age of human gene families (origin) was carried out using the ENSEMBL genome browser database. To identify the origin of individual human gene families, we fetched the phylogenies and analysed the evolutionary supertrees built by the pipeline of the ENSEMBL Compara multi-species comparisons project [52,53]. The used release (99) of the project contained 282 reference species including 277 vertebrata, 4 eumetazoa and 1 opisthokonta (*S. cerevisiae*) species. Note, that in these phylogenies the most ancient node can be the ancestor of yeast. The origin of the gene family was identified by taking the taxonomy level of the most ancient node of the phylogenetic supertrees. Taxonomy levels were broken into major nested age categories (Mammals, Vertebrates, Eumetazoa, Opisthokonta) similarly to previous studies [54].

To define the evolutionary origin of regions we built a customized pipeline that included collecting and mapping mutations from COSMIC database to ENSEMBL entries, constructing multiple sequence alignments of protein families and mapping the cancer regions among orthologs and paralogs. According to the ENSEMBL supertrees, protein sequences of human paralogs (including the cancer gene) and their orthologs were queried from the database using the Rest API function. Then, multiple sequence alignments of the corresponding sequences were created with MAFFT (default settings) [21]. Based on the sequence alignments, cancer regions were mapped onto the sequences. In the mapping step, cancer regions were considered as functional units (linear motifs, linkers, disordered domains) and borders of the regions were defined according to this. When the highly mutated regions covered only a single residue, it was extended to cover the known functional linear motif or using its sequence neighbourhood. Based on this, the subset of paralogs, in which the mapped cancer region was found to be conserved was identified.

Next, the set of sequences containing regions that showed evolutionary similarity to the mutated regions were identified among the collected orthologs and paralogs. Conservation of the regions among paralogs was evaluated relying on two strategies, by calculating the similarity of mutated positions in the cancer risk regions (see below) and based on HMM profiles. This consideration was taken into account in order to reduce the chance of false conservation interpretation arising from the difficulty of aligning disordered proteins. In the HMM profiles were built from conserved cancer regions of vertebrate model organisms using the HMMER (version 3.3) method [55]. The identified region hits were manually checked to minimize the chance of false positives or negatives. Next, we identified the evolutionary most distant relative in which the cancer region was declared to be conserved. As a result, the origin of the region could differ from the origin of the orthologous gene family, when paralogue sequences which contained the conserved motif had a more ancient origin. Basically, we treated the cancer risk regions as the founder of the family. The taxonomy level of this ortholog was defined as the level in which the cancer region emerged in the common ancestor of this ortholog and *H. sapiens*.

### Region conservation

Within the identified cancer risk region, some of the positions could be more heavily mutated and are likely to be more critical for the function of this region. We took this into account when calculating the region conservation. Mutations for each position collected from the COSMIC database were mapped to the corresponding ENSEMBL human entry. Based on the sequence alignment corresponding to the cancer risk regions, we identified the positions that were similar to the reference sequence. Two positions were considered similar when the substitution score was not negative according to the BLOSUM 62 substitution matrix. A given cancer region was considered to be conserved between homologs, when the conserved residues carried more than 50% of missense mutations.

### Positive selection: Selectome and MK test results

For each entry in our dataset we collected information about positive selection using the Selectome database (current version 6) [12]. This database contains collected sites of positive selection detected on a single branch of the phylogeny using the systematic branch-site test of the CODEML algorithm from the PAML [56] phylogenetic package version 4b. The ratio of non-synonymous and synonymous substitutions (ω) can be interpreted as a measurement of selective pressure indicating purifying (ω values < 1), neutral (ω values = 1) or positive (ω values > 1) selection. In our work, positions under positive selection that have a posterior probability higher than 0.9 were extracted from the database and mapped onto our gene set.

However, the branch-site model generally cannot detect species-specific positive selection. Potential cases of human-specific positive selection may be achieved effectively by comparing divergence to polymorphism data, as in the McDonald and Kreitman (MK) test. Human-specific positive selection detected by MK test previously calculated [13] were mapped onto our dataset disordered cancer genes.

## Supplementary Materials

### Evolutionary origin of selected cases

Basically, evolutionary origin is defined based on the most recent common ancestor of a given orthologous group. However, in some cases the correct estimation of evolutionary origin is not a trivial task neither at the region nor at the gene-family level, and significantly different results can be obtained depending on the applied approach. Here, we present the more challenging cases by pointing out the main source of ambiguities.

One typical scenario when largely different gene origin estimations can be obtained corresponds to large families with complex architecture. This happens for example in the case of kinases, which is represented by **CSF1R** in our dataset. On one end of the spectrum, the founder gene approach focuses on the evolutionary novelties associated with a particular function (i.e. enzymatic activity). Accordingly, it compiles every gene as an ortholog that has kinase domain from Archaea to vertebrata [4]. On the other end, other orthology prediction methods which rely on entire sequences, not only the kinase domain, estimate a more recent (Vertebrata) gene age [54].

The representation of opisthokonts by only *S. cerevisiae* can lead to false prediction of origin. For example, ENSEMBL Compara cannot detect any *S. cerevisiae* orthologs for **SRSF2.** However, in previous studies, the origin of this gene was defined consistently to be unicellular. The reason for this contradiction is that the classical SRSF homologous proteins are found only in the fungus *S. pombe*, but they are missing from *S. cerevisiae* [54,57].

High disorder content can also cause ambiguities in automated methods. An example that illustrates the issue with extensive disorder corresponds to **SETBP1**, which encodes a protein predicted to be disordered in almost 100%. Several previous studies using different age-estimation approaches, including the founder-gene approach, the origin of this case was predicted to be the time of emergence of vertebrates [4,54]. However, based on the ENSEMBL Compara database, we predict a more ancient, Opisthokonta emergence.

Estimation of evolutionary origin can also be complicated by gene loss and depend on the availability of genome sequences. The evolutionary origin of **CBL** was predicted previously to be the emergence time of opisthokonts [58], however our results show an eumetazoa origin. On the one hand, the CBL protein family indeed has ancient origin with a predicted ortholog from Dictyostelium, which in fact means a more ancient, Eukaryota origin. On the other hand, results show that CBL ortholog was not identified in opisthokonts, meaning a putative evolutionary gap in the nested age-estimation system. As a result, in our work an Eumetazoa origin was retrieved. Furthermore, in terms of the cancer region, results show that Dictyostelium-CBL does not appear to have the helical linker region, where the cancer mutations occur [59,60].

Similarly, the phylogeny of **BCL2** is also unclear beyond eumetazoa [61–66]. In addition, this example is also a unique case in terms of region conservation. The BH4 family of the cancer region is clearly conserved in all vertebrates, however, the C-terminal linker region shows a different conservation pattern. The extension and sequence homology of this segment is variant among vertebrates, only mammalian orthologs show a clear similarity. Beyond that significant differences can be observed both in terms of region length and homology (Figure S1).

**Figure S1.**
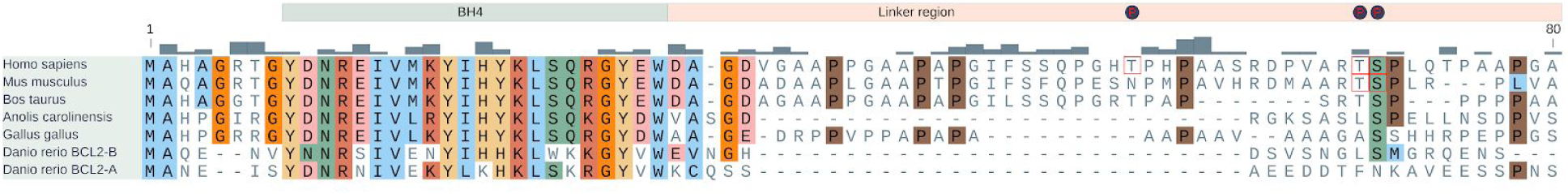
Sequence alignment of BCL2 cancer region. BH4 family is depicted by a light green box at the top of the alignment. Distribution of missense mutation is shown by gray bars. Dark circles and red rectangles show the positions of experimentally proven phosphorylation sites.

**Figure S2.**
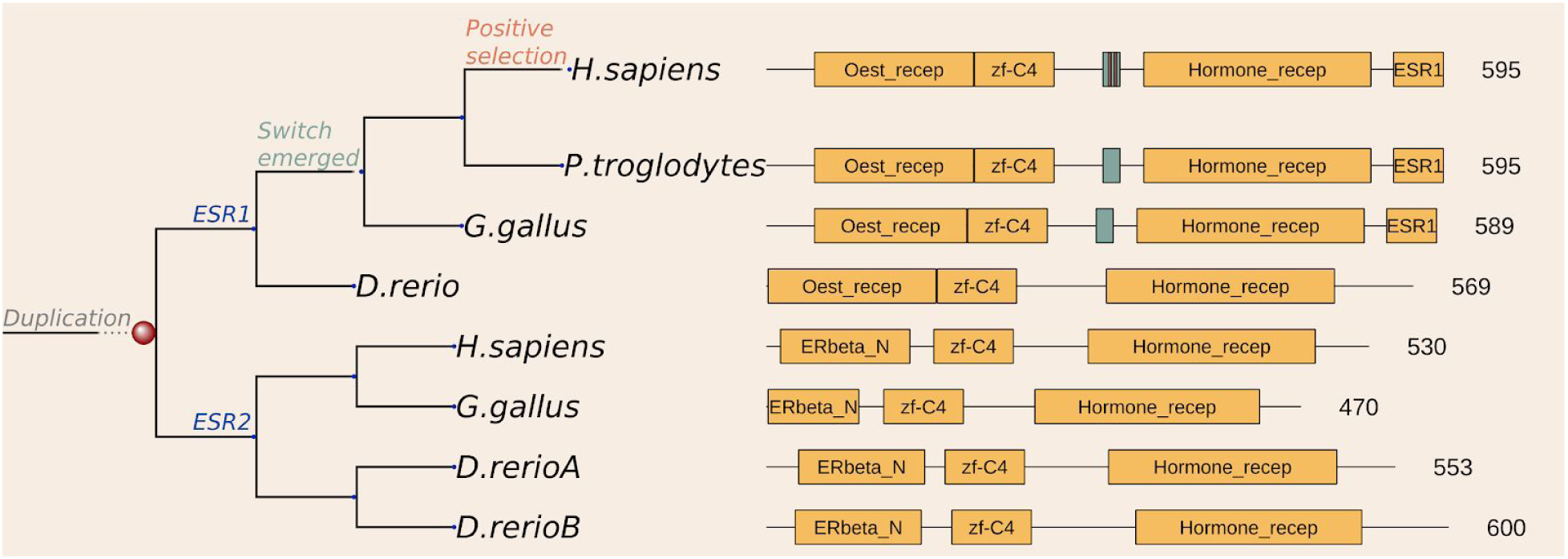
Schematic representation of the evolutionary scenario and functional units of the ESR1 and ESR2 proteins. Switch regions and domains are depicted by green and yellow boxes, respectively. Red stripes in the switch region of the human ESR1 indicate the positions of the *H. sapiens* specific positive selection.

## References

1. Jacob F. Evolution and tinkering. Science. 1977;196:1161–6.

2. Kinzler KW, Vogelstein B. Cancer-susceptibility genes. Gatekeepers and caretakers. Nature. 1997. p. 761, 763.

3. Domazet-Loso T, Tautz D. Phylostratigraphic tracking of cancer genes suggests a link to the emergence of multicellularity in metazoa. BMC Biol. 2010;8:66.

4. Domazet-Loso T, Tautz D. An ancient evolutionary origin of genes associated with human genetic diseases. Mol Biol Evol. 2008;25:2699–707.

5. Dickerson JE, Robertson DL. On the origins of Mendelian disease genes in man: the impact of gene duplication. Mol Biol Evol. 2012;29:61–9.

6. Bailey MH, Tokheim C, Porta-Pardo E, Sengupta S, Bertrand D, Weerasinghe A, et al. Comprehensive Characterization of Cancer Driver Genes and Mutations. Cell. 2018;173:371–85.e18.

7. Mészáros B, Zeke A, Reményi A, Simon I, Dosztányi Z. Systematic analysis of somatic mutations driving cancer: uncovering functional protein regions in disease development. Biol Direct. 2016;11:23.

8. Mészáros B, Hajdu-Soltész B, Zeke A, Dosztányi Z. Intrinsically disordered protein mutations can drive cancer and their targeted interference extends therapeutic options. Bioinformatics. bioRxiv; 2020. p. 2443.

9. Brown CJ, Johnson AK, Dunker AK, Daughdrill GW. Evolution and disorder. Curr Opin Struct Biol. 2011;21:441–6.

10. Davey NE, Cyert MS, Moses AM. Short linear motifs - ex nihilo evolution of protein regulation. Cell Commun Signal. 2015;13:43.

11. van den Berg TK, Yoder JA, Litman GW. On the origins of adaptive immunity: innate immune receptors join the tale. Trends Immunol. 2004;25:11–6.

12. Moretti S, Laurenczy B, Gharib WH, Castella B, Kuzniar A, Schabauer H, et al. Selectome update: quality control and computational improvements to a database of positive selection. Nucleic Acids Res. 2014;42:D917–21.

13. Gayà-Vidal M, Albà MM. Uncovering adaptive evolution in the human lineage. BMC Genomics. 2014;15:599.

14. Wu G, Xu G, Schulman BA, Jeffrey PD, Harper JW, Pavletich NP. Structure of a beta-TrCP1-Skp1-beta-catenin complex: destruction motif binding and lysine specificity of the SCF(beta-TrCP1) ubiquitin ligase. Mol Cell. 2003;11:1445–56.

15. Shcherbakova PV, Kunkel TA. Mutator phenotypes conferred by MLH1 overexpression and by heterozygosity for mlh1 mutations. Mol Cell Biol. 1999;19:3177–83.

16. Takahashi M, Shimodaira H, Andreutti-Zaugg C, Iggo R, Kolodner RD, Ishioka C. Functional analysis of human MLH1 variants using yeast and in vitro mismatch repair assays. Cancer Res. 2007;67:4595–604.

17. Ohsawa T, Sahara T, Muramatsu S, Nishimura Y, Yathuoka T, Tanaka Y, et al. Colorectal cancer susceptibility associated with the hMLH1 V384D variant. Mol Med Rep. 2009;2:887–91.

18. Lee SE, Lee HS, Kim K-Y, Park J-H, Roh H, Park HY, et al. High prevalence of the MLH1 V384D germline mutation in patients with HER2-positive luminal B breast cancer. Sci Rep. 2019;9:10966.

19. Gueneau E, Dherin C, Legrand P, Tellier-Lebegue C, Gilquin B, Bonnesoeur P, et al. Structure of the MutLα C-terminal domain reveals how Mlh1 contributes to Pms1 endonuclease site. Nat Struct Mol Biol. 2013;20:461–8.

20. Kim Y, Furman CM, Manhart CM, Alani E, Finkelstein IJ. Intrinsically disordered regions regulate both catalytic and non-catalytic activities of the MutLα mismatch repair complex. Nucleic Acids Res. 2019;47:1823–35.

21. Katoh K, Standley DM. MAFFT multiple sequence alignment software version 7: improvements in performance and usability. Mol Biol Evol. 2013;30:772–80.

22. Kamura T, Maenaka K, Kotoshiba S, Matsumoto M, Kohda D, Conaway RC, et al. VHL-box and SOCS-box domains determine binding specificity for Cul2-Rbx1 and Cul5-Rbx2 modules of ubiquitin ligases. Genes Dev. 2004;18:3055–65.

23. Cardote TAF, Gadd MS, Ciulli A. Crystal Structure of the Cul2-Rbx1-EloBC-VHL Ubiquitin Ligase Complex. Structure. 2017;25:901–11.e3.

24. Sutovsky H, Gazit E. The von Hippel-Lindau tumor suppressor protein is a molten globule under native conditions: implications for its physiological activities. J Biol Chem. 2004;279:17190–6.

25. Cai Q, Verma SC, Kumar P, Ma M, Robertson ES. Hypoxia inactivates the VHL tumor suppressor through PIASy-mediated SUMO modification. PLoS One. 2010;5:e9720.

26. Cai Q, Robertson ES. Ubiquitin/SUMO modification regulates VHL protein stability and nucleocytoplasmic localization. PLoS One [Internet]. 2010;5. Available from: http://dx.doi.org/10.1371/journal.pone.0012636

27. Minervini G, Mazzotta GM, Masiero A, Sartori E, Corrà S, Potenza E, et al. Isoform-specific interactions of the von Hippel-Lindau tumor suppressor protein. Sci Rep. 2015;5:12605.

28. German P, Bai S, Liu X-D, Sun M, Zhou L, Kalra S, et al. Phosphorylation-dependent cleavage regulates von Hippel Lindau proteostasis and function. Oncogene. 2016;35:4973–80.

29. Qi H, Gervais ML, Li W, DeCaprio JA, Challis JRG, Ohh M. Molecular cloning and characterization of the von Hippel-Lindau-like protein. Mol Cancer Res. 2004;2:43–52.

30. Dhayalan A, Kudithipudi S, Rathert P, Jeltsch A. Specificity analysis-based identification of new methylation targets of the SET7/9 protein lysine methyltransferase. Chem Biol. 2011;18:111–20.

31. Wang C, Fu M, Angeletti RH, Siconolfi-Baez L, Reutens AT, Albanese C, et al. Direct acetylation of the estrogen receptor alpha hinge region by p300 regulates transactivation and hormone sensitivity. J Biol Chem. 2001;276:18375–83.

32. Wang R-A, Mazumdar A, Vadlamudi RK, Kumar R. P21-activated kinase-1 phosphorylates and transactivates estrogen receptor-alpha and promotes hyperplasia in mammary epithelium. EMBO J. 2002;21:5437–47.

33. Michalides R, Griekspoor A, Balkenende A, Verwoerd D, Janssen L, Jalink K, et al. Tamoxifen resistance by a conformational arrest of the estrogen receptor alpha after PKA activation in breast cancer. Cancer Cell. 2004;5:597–605.

34. Cui Y, Zhang M, Pestell R, Curran EM, Welshons WV, Fuqua SAW. Phosphorylation of estrogen receptor alpha blocks its acetylation and regulates estrogen sensitivity. Cancer Res. 2004;64:9199–208.

35. Rust HL, Thompson PR. Kinase consensus sequences: a breeding ground for crosstalk. ACS Chem Biol. 2011;6:881–92.

36. de Leeuw R, Flach K, Bentin Toaldo C, Alexi X, Canisius S, Neefjes J, et al. PKA phosphorylation redirects ERα to promoters of a unique gene set to induce tamoxifen resistance. Oncogene. 2013;32:3543–51.

37. Atsriku C, Britton DJ, Held JM, Schilling B, Scott GK, Gibson BW, et al. Systematic mapping of posttranslational modifications in human estrogen receptor-alpha with emphasis on novel phosphorylation sites. Mol Cell Proteomics. 2009;8:467–80.

38. Williams CC, Basu A, El-Gharbawy A, Carrier LM, Smith CL, Rowan BG. Identification of four novel phosphorylation sites in estrogen receptor alpha: impact on receptor-dependent gene expression and phosphorylation by protein kinase CK2. BMC Biochem. 2009;10:36.

39. Walker VR, Korach KS. Estrogen receptor knockout mice as a model for endocrine research. ILAR J. 2004;45:455–61.

40. Porteous R, Herbison AE. Genetic Deletion of Esr1 in the Mouse Preoptic Area Disrupts the LH Surge and Estrous Cyclicity. Endocrinology. 2019;160:1821–9.

41. van der Lee R, Buljan M, Lang B, Weatheritt RJ, Daughdrill GW, Dunker AK, et al. Classification of intrinsically disordered regions and proteins. Chem Rev. 2014;114:6589–631.

42. Lengauer C, Kinzler KW, Vogelstein B. Genetic instabilities in human cancers. Nature. 1998;396:643–9.

43. Ellison AR, Lofing J, Bitter GA. Human MutL homolog (MLH1) function in DNA mismatch repair: a prospective screen for missense mutations in the ATPase domain. Nucleic Acids Res. 2004;32:5321–38.

44. Duchatel RJ, Jackson ER, Alvaro F, Nixon B, Hondermarck H, Dun MD. Signal Transduction in Diffuse Intrinsic Pontine Glioma. Proteomics. 2019;19:e1800479.

45. Piazza R, Magistroni V, Redaelli S, Mauri M, Massimino L, Sessa A, et al. SETBP1 induces transcription of a network of development genes by acting as an epigenetic hub. Nat Commun. 2018;9:2192.

46. Martin-Marcos P, Zhou F, Karunasiri C, Zhang F, Dong J, Nanda J, et al. eIF1A residues implicated in cancer stabilize translation preinitiation complexes and favor suboptimal initiation sites in yeast. Elife [Internet]. 2017;6. Available from: http://dx.doi.org/10.7554/eLife.31250

47. Bretones G, Álvarez MG, Arango JR, Rodríguez D, Nadeu F, Prado MA, et al. Altered patterns of global protein synthesis and translational fidelity in RPS15-mutated chronic lymphocytic leukemia. Blood. 2018;132:2375–88.

48. Masaki S, Ikeda S, Hata A, Shiozawa Y, Kon A, Ogawa S, et al. Myelodysplastic Syndrome-Associated SRSF2 Mutations Cause Splicing Changes by Altering Binding Motif Sequences. Front Genet. 2019;10:338.

49. Assis R, Bachtrog D. Rapid divergence and diversification of mammalian duplicate gene functions. BMC Evol Biol. 2015;15:138.

50. Trigos AS, Pearson RB, Papenfuss AT, Goode DL. How the evolution of multicellularity set the stage for cancer. Br J Cancer. 2018;118:145–52.

51. Sondka Z, Bamford S, Cole CG, Ward SA, Dunham I, Forbes SA. The COSMIC Cancer Gene Census: describing genetic dysfunction across all human cancers. Nat Rev Cancer. 2018;18:696–705.

52. Yates AD, Achuthan P, Akanni W, Allen J, Allen J, Alvarez-Jarreta J, et al. Ensembl 2020. Nucleic Acids Res. 2020;48:D682–8.

53. Herrero J, Muffato M, Beal K, Fitzgerald S, Gordon L, Pignatelli M, et al. Ensembl comparative genomics resources. Database [Internet]. 2016;2016. Available from: http://dx.doi.org/10.1093/database/bav096

54. Liebeskind BJ, McWhite CD, Marcotte EM. Towards Consensus Gene Ages. Genome Biol Evol. 2016;8:1812–23.

55. Eddy SR. Accelerated Profile HMM Searches. PLoS Comput Biol. 2011;7:e1002195.

56. Yang Z. PAML 4: phylogenetic analysis by maximum likelihood. Mol Biol Evol. 2007;24:1586–91.

57. Busch A, Hertel KJ. Evolution of SR protein and hnRNP splicing regulatory factors. Wiley Interdiscip Rev RNA. 2012;3:1–12.

58. Chu X-Y, Jiang L-H, Zhou X-H, Cui Z-J, Zhang H-Y. Evolutionary Origins of Cancer Driver Genes and Implications for Cancer Prognosis. Genes [Internet]. 2017;8. Available from: http://dx.doi.org/10.3390/genes8070182

59. Mohapatra B, Ahmad G, Nadeau S, Zutshi N, An W, Scheffe S, et al. Protein tyrosine kinase regulation by ubiquitination: critical roles of Cbl-family ubiquitin ligases. Biochim Biophys Acta. 2013;1833:122–39.

60. Langenick J, Araki T, Yamada Y, Williams JG. A Dictyostelium homologue of the metazoan Cbl proteins regulates STAT signalling. J Cell Sci. 2008;121:3524–30.

61. Lanave C, Santamaria M, Saccone C. Comparative genomics: the evolutionary history of the Bcl-2 family. Gene. 2004;333:71–9.

62. Hsu SY, Kaipia A, McGee E, Lomeli M, Hsueh AJ. Bok is a pro-apoptotic Bcl-2 protein with restricted expression in reproductive tissues and heterodimerizes with selective anti-apoptotic Bcl-2 family members. Proc Natl Acad Sci U S A. 1997;94:12401–6.

63. Kawai M, Pan L, Reed JC, Uchimiya H. Evolutionally conserved plant homologue of the Bax inhibitor-1 (BI-1) gene capable of suppressing Bax-induced cell death in yeast(1). FEBS Lett. 1999;464:143–7.

64. Greenhalf W, Stephan C, Chaudhuri B. Role of mitochondria and C-terminal membrane anchor of Bcl-2 in Bax induced growth arrest and mortality in Saccharomyces cerevisiae. FEBS Lett. 1996;380:169–75.

65. Banjara S, Suraweera CD, Hinds MG, Kvansakul M. The Bcl-2 Family: Ancient Origins, Conserved Structures, and Divergent Mechanisms. Biomolecules [Internet]. 2020;10. Available from: http://dx.doi.org/10.3390/biom10010128

66. Strasser A, Vaux DL. Viewing BCL2 and cell death control from an evolutionary perspective. Cell Death Differ. 2018;25:13–20.

